# Single nucleus sequencing fails to detect microglial activation in human tissue

**DOI:** 10.1101/2020.04.13.035386

**Authors:** N. Thrupp, C. Sala Frigerio, L. Wolfs, N. G. Skene, S. Poovathingal, Y. Fourne, P. M. Matthews, T. Theys, R. Mancuso, B. de Strooper, M. Fiers

## Abstract

Single nucleus RNA-Seq (snRNA-Seq) methods are used as an alternative to single cell RNA-Seq methods, as they allow transcriptomic profiling of frozen tissue. However, it is unclear whether snRNA-Seq is able to detect cellular state in human tissue. Indeed, snRNA-Seq analyses of human brain samples have failed to detect a consistent microglial activation signature in Alzheimer’s Disease. A comparison of microglia from single cells and single nuclei of four human subjects reveals that ~1% of genes is depleted in nuclei compared to whole cells. This small population contains 18% of genes previously implicated in microglial activation, including *APOE, CST3, FTL, SPP1*, and *CD74*. We confirm our findings across multiple previous single nucleus and single cell studies. Given the low sensitivity of snRNA-Seq to this population of activation genes, we conclude that snRNA-Seq is not suited to detecting cellular activation in microglia in human disease.

## Introduction

Single cell approaches allow us to study cell-to-cell heterogeneity (Habib et al., 2017), in brain material however, it is difficult to dissociate individual cells (Habib et al., 2017; Lake et al., 2016). This is further complicated if one is interested in studying the human brain, where often only frozen material is available. One alternative to study cellular transcriptional heterogeneity in brain tissue is single nucleus transcriptomics. Single nucleus RNA-Seq (snRNA-Seq) studies have shown concordance between single cell and single nucleus transcriptome profiles in mice (Bakken et al., 2018; Habib et al., 2017; Lake et al., 2017), but have limited the comparison to the identification of major cell types. It is unclear whether a snRNA-Seq approach is equally effective in identifying dynamic cellular substates such as microglial activation in human tissue.

A recent breakthrough in the field of Alzheimer’s Disease (AD) using single cell RNA-Seq (scRNA-Seq) demonstrated clearly that microglia become activated in response to amyloid plaques in mouse models (Keren-Shaul et al., 2017). This response comprises a transcriptional switch to a state called Activation Response Microglia (ARM) (Sala Frigerio et al., 2019), or Disease-Associated Microglia (DAM, MGnD) (Keren-Shaul et al., 2017; Krasemann et al., 2017). Ample evidence suggests that this microglial response is also relevant in human AD: microglia are believed to play a role in amyloid clearance (Efthymiou and Goate, 2017) and complement-mediated synapse loss (Fonseca et al., 2017), and histological studies have demonstrated considerable microgliosis around plaques in humans (McGeer et al., 1987). In addition, there is significant overlap between those genes involved in the microglial response, and genes within loci carrying AD genetic risk, as identified in Genome-Wide Association Studies (GWAS) (Efthymiou and Goate, 2017; Jansen et al., 2019; Kunkle et al., 2019; Lambert et al., 2013; Marioni et al., 2018), for example, *APOE*, *TREM2*, *APOC1*, *CD33* (Sala Frigerio et al., 2019). Most recently, the engrafting of human microglia into AD mouse models, followed by single cell RNA-sequencing, identified 66 DAM genes relevant to human activation^15^, and a bulk RNA-Seq study of AD patients identified 64 DAM genes^16^. In stark contrast, a number of high-profile snRNA-Seq studies of microglia in human AD (Del-Aguila et al., 2019; Grubman et al., 2019; Mathys et al., 2019; Zhou et al., 2020) have not recovered a consistent microglial activation signature. A recent cluster analysis by Mathys *et al.* of 48 AD patients and controls reported only 28 of 257 orthologous activation genes in common with the DAM signature (Mathys et al., 2019). Differential expression analysis between AD and control patients revealed 22 genes upregulated in AD patients (5 overlapping with the DAM signature). Of these AD genes, only 8 were also upregulated in another snRNA-Seq study of human AD (Grubman et al., 2019), and only 4 were also upregulated in another snRNA-Seq study of AD TREM2 variants (Zhou et al., 2020). The AD TREM2 variant study also only identified 11 DAM genes enriched in AD patients compared with controls. Del Aguila *et al.*, analysing single nucleus transcriptomics from 3 AD patients, were unable to recapitulate an activation signature (Del-Aguila et al., 2019). This has led to speculation that there is no such DAM signature in humans.

Here we compared the performance of snRNA-Seq to scRNA-Seq for the analysis of microglia from human cortical biopsies, and demonstrated that technical limitations inherent to snRNA-Seq provide a more likely explanation for this lack of consistency in snRNA-Seq studies of AD. We confirmed our results using publicly-available data.

## Results

### snRNA-Seq recovers major cell types from human tissue, but not microglial state

scRNA-Seq of FACS-sorted microglia was performed on temporal cortices of four human subjects who had undergone neocortical resection (see Supplementary Table 1 for subject data)(Mancuso et al., 2019). We generated snRNA-Seq libraries from these same subjects. Following quality filtering, PCA analysis and clustering of 37,060 nuclei, we identified 7 major cell types (Supplementary Fig. 1a, b): oligodendrocytes (ODC, 34.0%), excitatory neurons (27.0%), interneurons (11.2%), oligodendrocyte precursors (OPC 9,4%), microglia (11.3%), astrocytes (6.0%), and endothelial cells (1.1%). We focus here on the microglial population, which was extracted from the main dataset.

We first checked whether clustering analysis of single nuclei could recover subpopulations of microglia comparable to the single cell approach. A comparison of single nucleus and single cell clustering suggested that we could only partially recover similar microglial subcluster structure using both methods (see Supplementary Text and Supplementary Fig. 1c-e).

### Gene expression profiling of human nuclei and cells

To compare gene abundance in single microglial cells (14,823 cells) and nuclei (3,940 nuclei), we performed a differential abundance analysis between cells and nuclei from the 4 subjects (Fig. 1a). As demonstrated in previous studies (Bakken et al., 2018; Gerrits et al., 2019; Habib et al., 2017; Lake et al., 2017), the majority of genes showed similar normalized abundance levels in cells and nuclei, with 98.6% of genes falling along the diagonal in Fig. 1a (Pearson’s correlation coefficient = 0.92, p < 2.2e-16). However, we identified a group of 246 genes (1.1% of detected genes) that was less abundant in nuclei (fold change < −2, p_adj_ < 0.05, blue in Fig. 1a). A second population of 68 genes (0.3%) was found to be more abundant in nuclei (fold change > 2, p_adj_ < 0.05, red in Fig. 1a). Additionally, 3,248 genes were exclusively detected in cells, and 5,068 genes exclusively detected in nuclei.

**Fig. 1:**
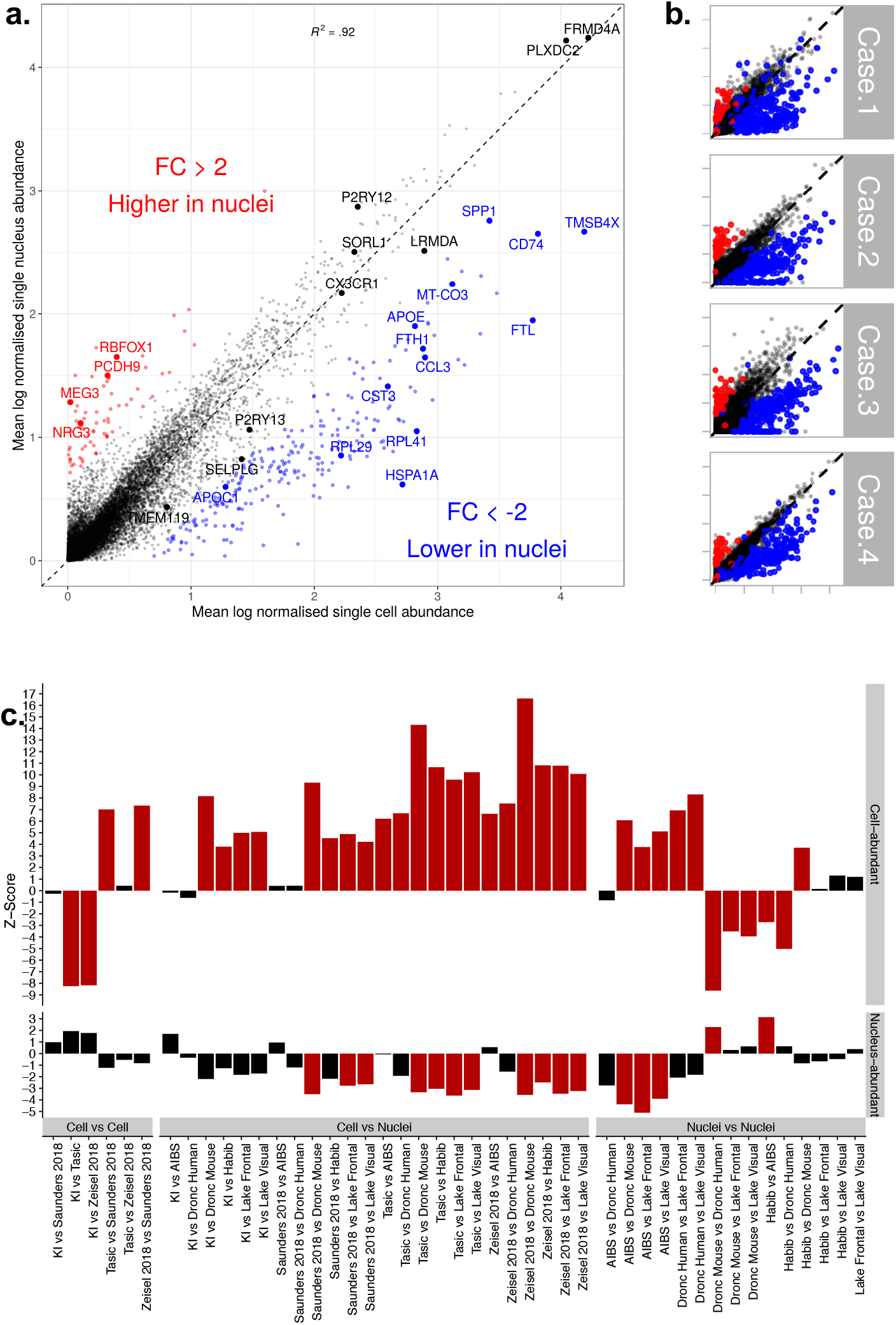
Gene abundance in single microglial cells versus single microglial nuclei of human cortical tissue. **a.** Mean normalised gene abundance in cells (x axis) and nuclei. (y axis). Red: genes with significantly higher abundance in nuclei (p_adj_ < 0.05, fold change > 2). Blue: genes that are significantly less abundance in nuclei (p_adj_ < 0.05, fold change < −2). Genes were normalized to read depth (per cell), scaled by 10,000 and log-transformed. *MALAT1* (which had normalized abundance levels of 6.0 and 6.9 respectively in cells and nuclei) has been removed for visualisation purposes. FC = fold change. Full results are available in Supplementary Table 2. **b.** Scatterplot as in a), per patient (with the same genes highlighted). Supplementary Table 1 contains patient data. **c.** Each bar represents a comparison between two datasets (*X* versus *Y*), with the bootstrapped *z*-scores representing the extent to which cell-enriched genes (top panel) and nuclear-enriched genes (bottom panel) have lower specificity for microglia in dataset *Y* relative to that in dataset *X*. Larger *z*-scores indicate greater depletion of genes, and red bars indicate a statistically significant depletion (p._adj_ < 0.05, by bootstrapping). KI = Karolinska Institutet; AIBS = Allen Institute for Brain Science.

The observed differences in abundance between cells and nuclei were consistent across all four subjects (Fig. 1b, Supplementary Fig. 2a). Downsampling of cellular reads indicated that differences in abundance were not the result of different sequencing depths (Supplementary Fig. 2b,c). The full differential abundance results can be found in Supplementary Table 2.

To assess the robustness of this finding, we used our nuclei-abundant genes and cell-abundant genes to compare enrichment across all pairs of 8 publicly-available single cell or single nucleus datasets (Supplementary Table 3, Fig. 1c). We consistently found our nuclei-depleted (cell-abundant) genes to be depleted in other single nucleus microglia compared to single cell microglia (mean microglial Z-score of cell-abundant genes was 7.95 when comparing cells to nuclei, whereas cell-to-cell comparisons yielded a mean of 0.01, and nuclei-to-nuclei comparisons yielded a mean of 0.81, for Z-scores with p_adj_ < 0.05). We also found our nuclei-abundant genes to be consistently enriched in other microglial nuclei compared with microglial cells (mean microglial Z-score −2.99 compared to −2.33 in nuclei against nuclei, no significant enrichment was found in cell-to-cell comparisons with p_adj_ < 0.05).

To assess functional enrichment among genes found to be more abundant in cells or nuclei, we ranked all genes according to log fold change (genes with a low abundance in nuclei had a negative log fold change) and performed a Gene Set Enrichment Analysis (GSEA, Subramanian et al., 2005) against gene markers from previous studies (Fig. 2a). For these analyses, a positive Normalised Enrichment Score (NES) represented nuclear enrichment, and a negative NES represented nuclear depletion. As expected, cytoplasmic RNA (defined by Bahar Halpern et al., 2015) was clearly enriched among genes found to be more abundant in cells (NES = −1.98, p_adj_ = 3.6e-05), as was mitochondrial mRNA (NES = −1.71, p_adj_ = 1.6e-04, gene set extracted from Ensembl’s BioMart (Zerbino et al., 2018)). mRNA found to be more abundant in the nucleus by (Bahar Halpern et al., 2015) tended towards enrichment in nuclei but was not significant (NES = 0.87, p_adj_ = 8.2e-01), which is to be expected as scRNA-Seq captures both nuclear and cytoplasmic RNA. RNA of genes coding for ribosomal proteins were also depleted in nuclei (NES = −2.28, p_adj_ = 3.6e-05), as previously described^1^. Genes with shorter coding sequences (CDS) were depleted in nuclei (NES = −1.38, p_adj_ = 2.5e-02), while longer CDS were enriched (NES = 2.07, p_adj_ = 2.1e-05), as already observed in earlier snRNA-Seq studies^3^. Finally, the genes defined by Gerrits (Gerrits et al., 2019) as cellular-enriched in a differential analysis of microglial cells versus (fresh) nuclei in humans were also enriched in cells in our data, showing a NES score of −2.15 (p_adj_ = 3.6e-05).

**Fig. 2:**
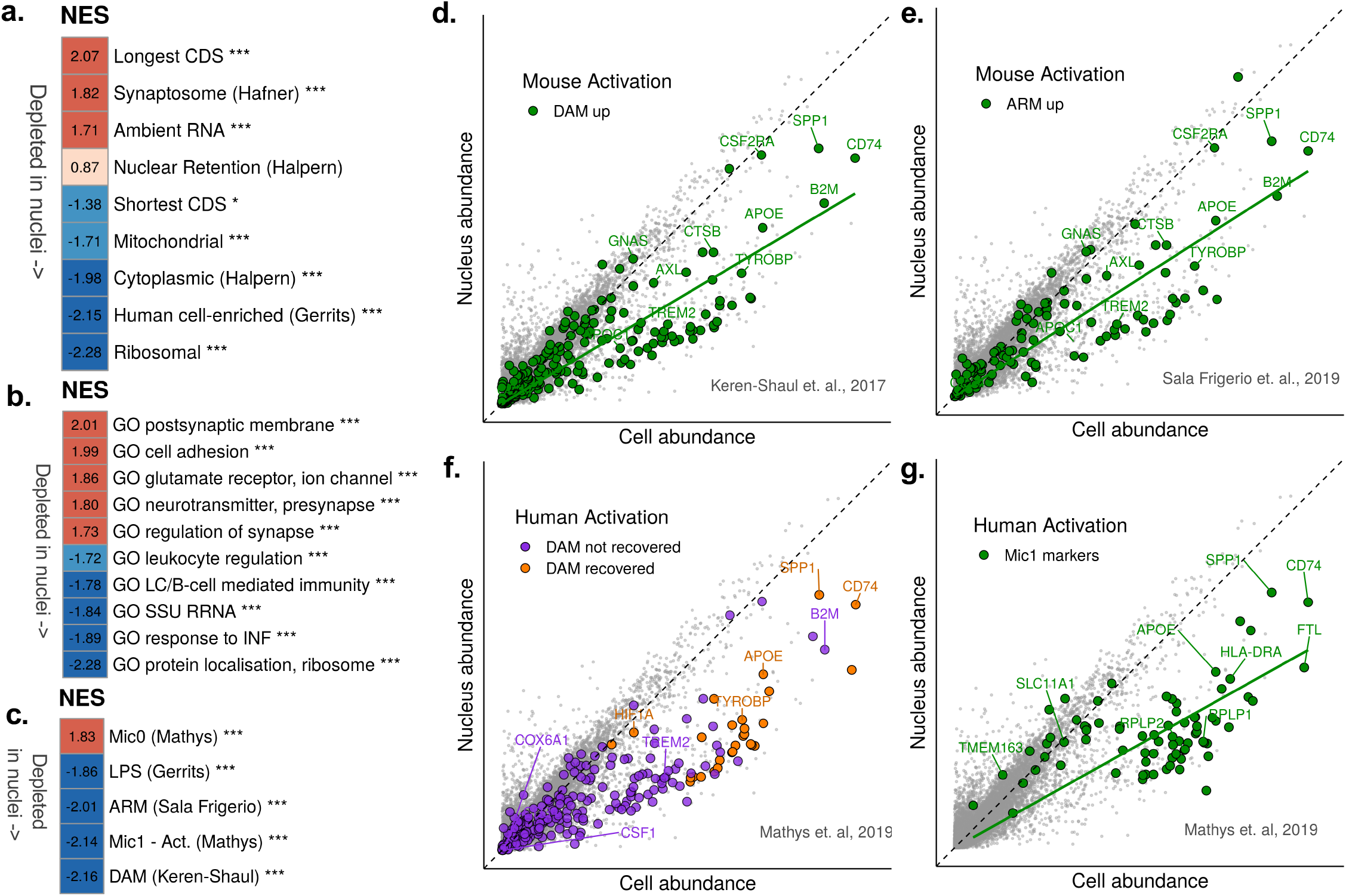
Functional analysis of genes that are enriched or depleted in nuclei. **a.** Gene Set Enrichment Analysis (GSEA) of gene sets related to cellular location and gene coding sequence length (CDS). Background genes were ranked according to log fold change of nuclei versus cells. Red: higher Normalised Enrichment Score (NES), i.e. more genes associated with nuclear enrichment; blue: negative NES scores (depletion in nuclei). *** represents significance (p_adj_ < 0.0005). CDS = coding sequence. **b.** GSEA of *super*-Gene Ontology gene sets against ranked nucleus-cell log fold changes. Only top and bottom categories (according to NES) are shown. Colours as in a). GO = Gene Ontology; SSU RRNA = small subunit ribosomal RNA; INF = Interferon; LC = leukocyte. **c.** GSEA of selected gene sets from previous studies of microglial activation, against fold change as in a). *** represents significance (p_adj_ < 0.0005). Mic0 = markers of microglial cluster 0 in human brain tissue; Mic1 = markers of microglial cluster 1 (response to plaques) defined by (Mathys et al., 2019) in human brain tissue. ARM = Activation Response Microglia (Sala Frigerio *et al.*^8^). DAM = Disease-Associated Microglia (Keren-Shaul *et al.*^7^) **d.** Scatterplot as in Fig. 1a), highlighting in green the DAM genes. A regression line for the highlighted genes is shown in green (slope = 0.60). **e.** Scatterplot as in d), highlighting in green the ARM genes. A regression line for the highlighted genes is shown in green (slope = 0.64). **f.** Scatterplot as in d), highlighting the DAM genes recovered in the study of human activation in AD (Mathys et al., 2019). Purple: DAM genes not recovered in their study; orange: DAM genes recovered in their study. **g.** Scatterplot as in d), Green: human activation marker genes defined by (Mathys et al., 2019). A regression line for the highlighted genes is shown in green (slope = 0.56). Gene sets are available in Supplementary Table 6.

To further characterise genes with higher or lower abundance in nuclei compared with cells, we performed GSEA, using Gene Ontology (GO) terms extracted from MSigDb (Liberzon et al., 2011) against the ranked log fold change. We selected the 100 terms with the highest NES, and the 100 terms with the lowest NES (p_adj_ < 0.05). Given the high overlap in terms, we clustered ontology terms based on the number of shared genes, in order to define “super” GO clusters (Supplementary Fig. 2e,f). We repeated the GSEA analysis using these *super*-GO clusters (Fig. 2b, Supplementary Tables 4 and 5) and observed an enrichment of neuronal and synaptic terms in nuclei-abundant genes (also shown in the red population in Fig.1a). We suspect a synaptosome contamination during centrifugation. This is supported by the enrichment of synaptosome genes (NES = 1.82, p_adj_ = 3.6e-05, Fig. 2a; Hafner et al., 2019) and ambient RNA – mRNA originating not from cells/nuclei but from free-floating transcripts in the solution (Macosko et al., 2015) – (NES = 1.71, p_adj_ = 9.2e-05, Fig. 2a, Supplementary Fig. 2d) within the nucleus-abundant genes. The two gene sets share a strong overlap (Supplementary Table 6). These genes, although enriched in nuclei compared with cellular levels, still show low abundance (most of these genes show a normalised abundance of no more than 2 – Fig. 1a).

### Activation genes identified in mouse models of AD are depleted in human nuclei

More interesting was the depletion of immune-related genes in nuclei (Fig. 2b). We therefore tested whether microglial activation genes^7,8,17^ were also depleted in nuclei (Fig. 2c, Supplementary Tables 5 and 6). Remarkably, we found a strong depletion of genes associated with mouse microglial activation: 45 of 257 orthologous DAM genes (Keren-Shaul et al., 2017), (NES = −2.16, p_adj_ = 3.6e-05, Fig. 2c,d), and 28 of 200 orthologous ARM genes (Sala Frigerio et al., 2019) (NES = −2.01, p_adj_ = 3.6e-05, Fig. 2c,e), confirming that mouse microglial activation genes wereless abundant in nuclei. Genes upregulated by LPS stimulation in mice (Gerrits et al., 2019) also showed depletion in nuclei (NES = −1.86, p_adj_ = 3.6e-05, Fig. 2c, Supplementary Figure 2g).

### Activation genes identified in mouse studies of AD are depleted in human nuclei

We next examined genes that were identified as markers of the human microglial response to AD in the recent snRNA-Seq study by (Mathys et al., 2019) (Fig. 2c, f, g). Markers of this response (referred to by Mathys *et al.* as “Mic1”) had a NES score of −2.14 (p_adj_ = 3.6e-05), indicating that they were depleted in nuclei (Fig. 2c). The study identified 28 DAM genes as marker genes of the Mic1 response cluster (shown in orange in Fig. 2f); however the majority of DAM genes were not recovered using their snRNA-Seq protocol (purple in Fig. 2f). Fig. 2g shows in green all the markers of the human activation cluster Mic1. Clearly, DAM genes and other Mic1 markers showed higher abundance in cells relative to nuclei (confirming the NES score in Fig. 2c). Further, it seems likely that the recovered DAM genes (orange in Fig. 2f) and Mic1 markers in general (green in Fig. 2g), were detected in the original snRNA-Seq experiment owing to their higher nuclear abundance compared with the nuclear abundance of other genes, including those DAM genes that were not recovered (purple in Fig. 2f).

## Discussion

In summary, in our comparison of snRNA-Seq and scRNA-Seq of human microglia, we identified a set of genes (1.1% of the gene population) with at least 2-fold lower abundance in nuclei compared to their cellular levels (Fig. 1a-b). This small set is strongly enriched for genes previously associated with microglial activation in mouse models of AD, for example *APOE*, *CST3*, *FTL*, *SPP1*, and *CD74* (Fig. 2b-e). Thus, while our work agrees with previous experiments demonstrating that snRNA-Seq can determine cell type (Supplementary Fig. 1a,b), we argue that there are important limitations when studying cellular state in humans. This limitation is likely responsible for the difficulty in identifying consistent DAM- or ARM-like gene populations in the human brain in snRNA-Seq-based studies. We identified similar patterns of depletion in other single nucleus microglia (Fig. 2c).

Examination of data from the Mathys *et al.* study of human nuclei in AD (Mathys et al., 2019) shows that only genes with higher nuclear abundance levels were detected (Fig. 2f, g). This suggests that the discordance between human and mouse microglial activation is at least in part a consequence of limitations in the technology, rather than biological differences between the species as current snRNA-Seq suggest. Deeper sequencing (or increased sample sizes) may compensate for this lack of sensitivity. However, the sparse nature of snRNA-Seq and the high level of heterogeneity in human samples, combined with the fact that many relevant genes have a more than two-fold lower abundance in nuclei (*e.g. APOE* fold change = 2.57, *CST3* fold change = 3.44, *FTL* fold change = 6.53), strongly suggests that this will remain a problem.

While our data is (at least partially) in agreement (Fig. 2a, c) with Gerrits *et al.* (Gerrits et al., 2019) which also compares nuclei with cells in human microglia, they did not report a nuclear depletion of activation genes. We suspect the reason for this is (a) the low human sample number (n=2); (b) for the cluster analysis, Gerrits *et al.* scaled cell and nuclei expression to mitochondrial and ribosomal reads, essentially masking differences between nuclei and cells, and (c) for the differential expression analysis, Gerrits *et al.* compared fresh cells to fresh nuclei, as opposed to frozen nuclei.

Alternative approaches may be more suitable for generating a brain atlas of human disease such as AD, particularly where we are limited to frozen material. *In situ* spatial transcriptomics (ST) negates issues related to tissue dissociation and cell or nucleus isolation (Ståhl et al., 2016), while at the same time retaining spatial information. This approach has recently been applied to examine transcriptomic changes and identify genes that are co-expressed across multiple cell types in the amyloid plaque niche of the mouse brain (Chen et al., 2019). In humans, a similar methodology was recently applied to identify pathway dysregulation and regional differences in cellular states of the postmortem spinal tissue of Amyotrophic Lateral Sclerosis (ALS) patients states (Maniatis et al., 2019). Its application to AD patients may shed light on transcriptomic changes occurring in microglia which localize near plaques, and may also provide insights into the crosstalk occurring between neighbouring cells.

In conclusion, while snRNA-Seq offers a viable alternative to scRNA-Seq for identification of cell types in tissue where cell dissociation is problematic, its utility for detecting cellular states in disease is limited.

## Supporting information

S1_Patient_metadata

S2_Expression_cells_nuclei

S3_Public_datasets

S4_GO_annotations

S5_GSEA

S6_Genesets

S7_Cluster_markers

## Data Availability

Sequencing data from single microglial cells is available on GEO (accession number GSE137444). Sequencing data from single nuclei will be made available on GEO.

## Code Availability

Analysis of previous datasets was performed using the EWCE package (Skene and Grant, 2016) for R and the MicroglialDepletion package (https://github.com/NathanSkene/MicroglialDepletion).

## Acknowledgements

Work in the De Strooper laboratory was supported by the European Union (ERC-834682 - CELLPHASE_AD), the Fonds voor Wetenschappelijk Onderzoek (FWO), KU Leuven, VIB, UK-DRI (Medical Research council, ARUK and Alzheimer Society), a Methusalem grant from KU Leuven and the Flemish Government, the “Geneeskundige Stichting Koningin Elisabeth”, Opening the Future campaign of the Leuven Universitair Fonds (LUF), the Belgian Alzheimer Research Foundation (SAO-FRA) and the Alzheimer’s Association USA. Bart De Strooper is holder of the Bax-Vanluffelen Chair for Alzheimer’s Disease. Cell sorting was performed at the KU Leuven FACS core facility, and sequencing was carried out by the VIB Nucleomics Core. Renzo Mancuso is recipient of a postdoctoral fellowship from the Alzheimer’s Association, USA. Paul M. Matthews acknowledges generous support from the Edmond J. Safra Foundation and Lily Safra, the NIHR Biomedical Research Centre at Imperial College and the UK Dementia Research Institute.

## Author contributions

Conceptualization: B.D.S., M.F., C.S.F., R.M.; surgery and extraction of patient tissue samples: T.T.; Investigation: R.M., L.W., S.P.; formal analysis: N.T., Y.F., M.F., N.G.S., P.M.M.; Writing – original draft: N.T., C.S.F., M.F.; writing – review & editing: B.D.S., R.M., N.G.S., P.M.M.; supervision: M.F., B.D.S.

## Materials and Methods

### Isolation of human primary microglial cells

Human primary microglial cells from the Mancuso *et. al* study (Mancuso et al., 2019) were used. Briefly, microglia were FAC-sorted from brain tissue samples resected from the lateral temporal neocortex of 4 epilepsy patients during neurosurgery. The full protocol is described in the original study. All procedures followed protocols approved by the local Ethical Committee (protocol number S61186). Sequencing was performed as described for the nuclei.

### Isolation of nuclei from human subjects

Nuclei from frozen biopsy tissue were isolated as follows: brain tissue was sliced on dry ice, then homogenized manually (15 gentle strokes) with 1mL ice-cold Homogenisation Buffer (HB) with 5μL Rnasin Plus. The homogenate was strained with a 70μm strainer and washed with 1.65mL to a final volume of 2.65mL. 2.65mL of Gradient medium was added (Vf = 5.3mL). To isolate the nuclei, the sample was added to a 4mL 29% cushion using a P1000, and the weight adjusted with HB. The sample was centrifuged in a SW41Ti rotor at 7,700 rpm for 30 minutes at 4°C. The supernatant was removed with a plastic Pasteur pipette, followed by removal of the lower supernatant with P200. Nuclei were resuspended in 200μL of resuspension buffer, transferred to a new tube, washed again with 100-200μL resuspension buffer, and pooled with the previous solution. Clumps were disrupted by pipetting with P200, then filtered through a Falcon tube with 0.35μm strainer. 9μL of sample was mixed with 1μL of propidium iodide (PI) stain, loaded onto a LUNA-FL slide and allowed to settle for 30 seconds. We viewed nuclei with the LUNA-FL Automated cell counter to check numbers and shape.

### Single nucleus sequencing

RNA sequencing was performed using the 10X Genomics Single Cell 3` Reagent Kit (v2) according to manufacturer protocols. cDNA libraries from fresh-frozen nuclei were sequenced on an Illumina HiSeq platform 4000. Supplementary Table 1 provides sequencing information per sample (for cells and nuclei).

### Single nucleus analysis

#### Alignment

Cellranger v2.1.1 was used to demultiplex and align sequencing output to a human reference genome (assembly hg38 build 95). We used a “pre-mRNA” database to align single nuclei to exons as well as introns (10x Genomics, n.d.). Following alignment, nuclei from one patient sample (RM101.1) were removed due to poor quality (low read and gene count). See Supplementary Table 1 for sample information. Unfiltered count matrices were used for downstream analysis.

#### Extraction of microglial nuclei

Data was processed using the Seurat v3.0.2 package (Butler et al., 2018; Stuart et al., 2019) in R v3.6.1. For each patient, the count matrix was filtered to exclude nuclei with fewer than 100 genes. Counts were depth-normalised, scaled by 10,000 and log-transformed. *FindVariableFeatures* was run using a variance-stabilising transformation (“vst”) to identify the 2,000 most variable genes in each sample. Data from the 4 patients was then integrated using Seurat’s *FindIntegrationAnchors* with default parameters, and *IntegrateData* using 40 principal components (PCs). The dataset consisted of 37,060 nuclei, with a mean read depth of 4,305 counts per nucleus, and 1,791 genes per nucleus. Integrated data was scaled (default Seurat parameters). We ran a Principal Components Analysis (PCA), then calculated Uniform Manifold Approximation and Projection (UMAP) embeddings using 40 PCs. We identified clusters using Seurat’s *FindNeighbours* and *FindClusters* functions, again using 40 PCs. Based on abundance of known celltype markers, we assigned each cluster to a cell type. Microglial clusters were identified using known markers including *P2RY12*, *CSF1R*, *CX3CR1*, and extracted for downstream analysis.

#### Pre-processing of microglial nuclei per patient

Microglia from each patient sample were analysed individually as described for all cell types above, with the following modifications: raw counts were filtered to remove genes and counts that were ± 3 standard deviations away from the median value. After normalization, doublets were identified using DoubletFinder v2.0.2 (McGinnis et al., 2019) using 40 PCs, assuming a 7.5% doublet rate. Following removal of doublets, filtering and Seurat normalization were performed again. Data from patients was then integrated and clusters were identified as above. We discarded small clusters than contained markers for microglia as well as other cell types. After pre-processing, 3,927 nuclei remained, with a mean count depth of 1,295 and a mean gene count of 879 genes per nucleus.

### Single cell analysis

Full details of single cell processing are available in Mancuso *et al.*(Mancuso et al., 2019). Only cells from the four patients included in the single nucleus study were used here.

### Comparisons of single cells and single nuclei

#### Cluster analysis (see supplementary text)

In order to identify microglial cell states in the nuclei data we calculated gene markers for each cluster using Seurat’s *FindMarkers* function, selecting only markers with a positive fold change. Gene markers for cell clusters were extracted from the original Mancuso *et al.* study. Markers for nuclei and cells are available in Supplementary Table 7. For the analysis, we kept the top 40 significant markers (p_adj_ < 0.05) based on log fold change for the nuclear clusters and cellular clusters. For each nucleus, we calculated the mean abundance levels of each cell cluster marker set against the aggregated abundance of random control gene sets, using Seurat’s *AddModuleScore* function. This gave us the MS40 score for each marker set. We performed two-sided Fisher’s Exact tests with Benjamini Hochberg corrections to determine the overlap of cell cluster markers with nuclear cluster markers (selecting the top 40 markers for each set), using the union of all genes in the cell and nuclei datasets as a background (p_adj_ < 0.05 was considered significant).

#### Differential Abundance

We discarded all non-microglial clusters (brain macrophages, neutrophils), leaving 3,721 nuclei and 14,435 cells. Differential abundance analysis was performed with the Seurat package, using a two-sided Wilcoxon rank sum test, with a Bonferroni correction for multiple testing. Genes with p_adj_ < 0.05 and fold change > |2| were considered significant. As Seurat applies a pseudocount of +1 to data before calculating log fold changes, a fold change of 2 corresponds to a log fold change of 0.63. Log fold changes calculated by Seurat were used for further analysis in gene set enrichment analysis.

#### Scatter plots

We calculated the mean of the normalized abundance levels for cells and for nuclei, and log-transformed these values.

#### Assessment of nuclear-enriched or cell-enriched gene sets in public scRNA-Seq and snRNA-Seq datasets

We followed the methodology described in (Skene et al., 2018): genes that were significantly more abundant in nuclei or more abundant in cells (see “Differential abundance” methodology above) were used, creating two gene sets. 8 public datasets were reduced to contain six major cell types: pyramidal neurons, interneurons, astrocytes, interneurons, microglia and oligodendrocyte precursors. Within each dataset, for each gene in our gene sets, we calculated a celltype specificity score using the EWCE R package (Github version committed July 29, 2019; Skene and Grant, 2016). For each pair of datasets, *X* and *Y*, we subtracted the mean microglial specificity score of *Y* from *X.* We then calculated the same scores for 10,000 random gene sets: the probability and z-score for the difference in specificity for the dendritic genes is calculated using these. Finally, the depletion z-score for each gene set was equal to: (mean subtracted microglial specificity score – bootstrapped mean) / (bootstrapped standard deviation). A large positive z-score thus indicated that the gene set was depleted in microglia of dataset *Y* relative to dataset *X*. Benjamini-Hochberg multiple testing corrections were applied.

#### Public datasets

For the Karolinska Institutet (KI) dataset (Skene et al., 2018), we used S1 pyramidal neurons. For the Zeisel 2018 dataset (Zeisel et al., 2018) we used all ACTE* cells as astrocytes, TEGLU* as pyramidal neurons, TEINH* as interneurons, OPC as oligodendrocyte precursors and MGL* as microglia. For the Saunders dataset (Saunders et al., 2018), we used all Neuron.Slc17a7 celltypes from the frontal cortex (FC), hippocampus (HC) or posterior cortex (PC) as pyramidal neurons; all Neuron.Gad1Gad2 cell types from FC, HC or PC as interneurons; Polydendrocye as OPCs; Astrocyte as astrocytes, and Microglia as microglia. The Lake datasets both came from a single publication (Lake et al., 2018) which had data from frontal cortex, visual cortex and cerebellum. The cerebellum data was not used here. Data from frontal and visual cortices were analyzed separately. All other datasets - Dronc Human (Habib et al., 2017), Dronc Mouse (Habib et al., 2017), Allen Institute for Brain Science (AIBS) (Hodge et al., 2019), Tasic (Tasic et al., 2016) and Habib (Habib et al., 2016) – were used as described previously (Skene et al., 2018). Supplementary Table 3 lists all datasets. An R package is available for the analysis at https://github.com/NathanSkene/MicroglialDepletion.

#### Functional analysis

We performed Gene Set Enrichment Analysis (GSEA) using the R package fgsea v1.8.0 (Sergushichev, 2016), using default parameters. Gene sets were mapped against a list of genes ranked according to fold change between cellular abundance and nuclear abundance. Gene ontology (GO) sets were obtained from MSigDB (Liberzon et al., 2011; Subramanian et al., 2005). Other gene sets were obtained from previous studies (see Supplementary Table 6). p_adj_ < 0.05 (Benjamini-Hochberg correction) was considered significant.

#### Clustering of gene ontology terms

GSEA of GO terms resulted in many functional categories with overlapping genes. In order to reduce this redundancy, the top and bottom 100 GO terms according to normalized enrichment score (with p_adj_ < 0.05) were clustered as follows: a Jaccard index (the size of the intersection of the two datasets, divided by the size of the union of the two datasets, multiplied by 100) of the overlapping genes was calculated between each significant GO set. The resulting similarity matrix was converted to a dissimilarity matrix, and hierarchical clustering was performed on the matrix. We selected a k value of 16 to group the GO terms based on the hierarchical clustering (see Supplementary Table 2). Gene sets were merged, and each new “super” GO was assigned an annotation manually. GSEA analysis was performed on these *super*-GO gene sets as described above.

#### Gene sets from previous studies

We extracted gene sets from previous studies for this analysis. A full list of gene sets is available in Supplementary Table 6. Where data was selected from mouse datasets, we converted the mouse gene to its human ortholog using R’s BioMaRt package v2.40.5 (Durinck et al., 2009), selecting only orthologs that displayed 1-to-1 orthology. For the ARM gene set we selected the top 200 ARM genes based on log fold change (Sala Frigerio et al., 2019). For the Gerrits human gene set, we took the union of all genes that showed significant differential abundance between cells and nuclei (microglia) from donor 1 and donor 2 (Gerrits et al., 2019). For the LPS gene set, we took the union of all genes significantly upregulated in LPS in cells and in nuclei (microglia) from the Gerrits study (Gerrits et al., 2019).

#### Downsampling of cell counts

To match cell sequencing depth to nucleus sequencing depth (see Supplementary Fig. 2b,c), we sampled without replacement the number of reads in the cells by a proportion of 0.32, using the *downsampleMatrix* function of the DropletUtils R package v1.4.3 (Griffiths et al., 2018; Lun et al., 2019). This resulted in a read depth of 1,304 compared with the original read depth of 3,979 reads per cell.

#### Definition of ambient RNA profile in nuclei

We extracted nuclei with less than 700 counts from the original unfiltered raw count matrix of all cell types (resulting in 2,414 nuclei with a mean read depth of 590), and summed the gene counts, under the assumption that these were empty drops rather than nuclei. We took the top 150 genes to represent the ambient RNA profile. The mean read depth of these genes in the empty drops was 121 reads per cell.

## Supplementary Data

### Text

Supplementary Text: Clustering of microglial cells and nuclei in human cortical tissue.

### Tables

Supplementary Table 1: Patient Metadata

Supplementary Table 2: Differential abundance nuclei versus cells

Supplementary Table 3: Public single cell / single nucleus datasets

Supplementary Table 4: Clustering of Gene Ontology (GO) terms

Supplementary Table 5: Results of GSEA analysis

Supplementary Table 6: Lists of all gene sets

Supplementary Table 7: Cluster markers for single nucleus cluster analysis

**Fig. S1:**
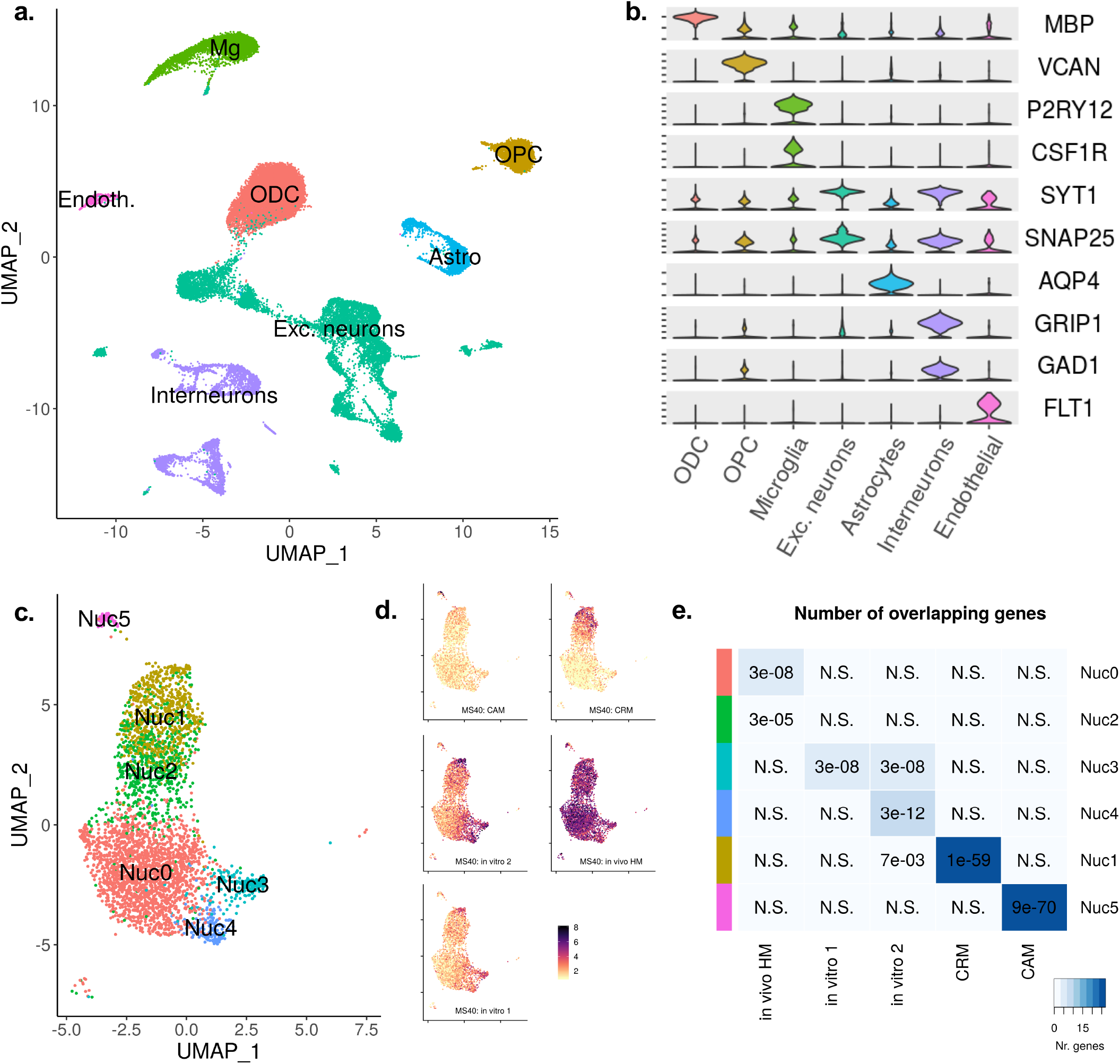
Clustering of single nuclei from human tissue. **a.** UMAP plot of 37,060 nuclei from cortical tissue of 4 neurosurgical patients, coloured according to cell type. Mg = Microglia; OPC = oligodendrocyte precursor cells; ODC = oligodendrocytes; Astro = Astrocytes; Endoth = Endothelial cells; Exc. neurons = excitatory neurons. **b.** Violin plots show selected markers of the different cell types (data is normalised for count depth and log-transformed). **c.** UMAP plot of 3,721 microglial nuclei from cortical tissue of 4 neurosurgical patients, coloured according to cluster number, after *in silico* extraction of microglia (based on markers such as *P2RY12*) and reclustering. **d.** Module scores for gene sets extracted from the original Mancuso *et al.* single cell microglia study (Mancuso et al., 2019). The top 40 genes according to log fold change were selected for each gene set. **e.** Overlap of top 40 marker genes from cellular clusters on the horizontal axis (Mancuso *et al.*) and nuclear clusters on the vertical axis. The blue scale represents the number of genes in common, numbers represent p_adj_ values. Vertical coloured bars correspond to the clusters shown in c). N.S. = not significant (p_adj_ > 0.05). MS40 = Module Score of top 40 gene markers; CAM = macrophages; CRM = cytokine response; in vitro 1 = activation-like module (similar to *in vitro* macrophages); in vitro 2 = activation-like module (similar to *in vitro* monocytes); in vivo HM = homeostatic. Nuc = Nuclear clusters. Cluster markers are provided in Supplementary Table 7.

**Fig. S2:**
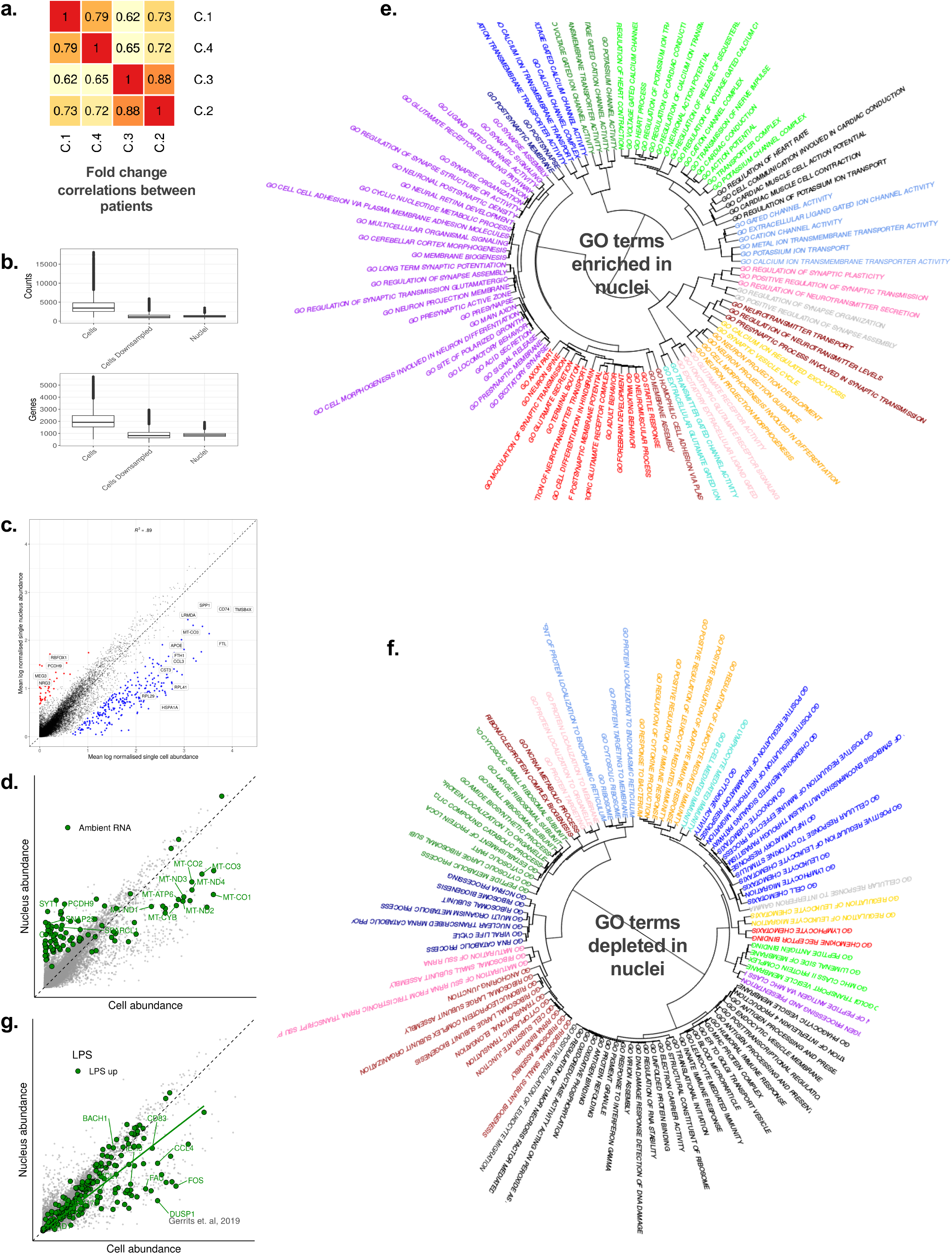
Gene abundance in single microglial cells versus single microglial nuclei of human cortical tissue. **a.** Correlation matrix of gene abundance fold changes (single cell vs single nucleus abundance) between patients. **b.** Downsampling of reads: boxplots for numbers of reads (top) and numbers of genes (bottom) for single cells before downsampling, single cells after downsampling, and single nuclei. Boxplots show median, with 25% and 75% quantiles. **c.** Scatterplot of mean gene abundance in cells against mean gene abundance in nuclei (as in Fig. 1a) after downsampling of reads in cells. Data is normalised to count depth and log-transformed. Points in red represent genes with significantly higher abundance in nuclei, while those in blue are significantly less abundant in nuclei (p_adj_ < 0.05, fold change > |2|). **d.** Scatter plot, as in Fig. 1a) showing the ambient mRNA in green (the same dataset was used in Fig. 2d). Ambient RNA is defined as the 150 most abundant genes in the 700 nuclei with the lowest total read counts. **e, f.** Dendrograms of **e.** top 100 Gene Ontology (GO) terms enriched in nuclei, and **f.** top 100 GO terms depleted in nuclei. GO terms were clustered based on overlap between their gene sets. The colours show how GO terms were clustered. These clusters are described in Supplementary Table 4. **g.** Scatterplot as in Fig. 1a), highlighting in green genes that are upregulated during LPS stimulation in mice (Gerrits et al., 2019). A regression line for the highlighted genes is shown in green (slope = 0.78).

### Supplementary Text: Clustering of microglial cells and nuclei in human cortical tissue

We sequenced nuclei from cortical tissue of 4 neurosurgical patients. Single cell sequencing of FAC-sorted microglia was performed on cortical tissue of the same patients in a previous study(Mancuso et al., 2019). Subject data is available in Supplementary Table 1. Following quality filtering, data integration, PCA analysis and clustering of nuclei, we identified 7 main cell types in 37,060 nuclei: oligodendrocytes (ODC, 34.0%), excitatory neurons (27.0%), interneurons (11.3%), oligodendrocyte precursors (OPC 9,4%), microglia (11.3%), astrocytes (6.0%), and endothelial cells (1.1%). Supplementary Fig. 1a and Supplementary Fig. 1b show UMAP embeddings for all nuclei, coloured by cell type, and selected markers for each cell type, respectively.

Microglial nuclei were isolated and reclustered. We identified 3,721 microglia (expressing *MEF2A*, *P2RY12*, *CX3CR1*, *CSF1R*), a macrophage cluster (enriched for *CD163* and *MRC1*, 67 nuclei), a neutrophil cluster (72 nuclei), and a cluster containing microglial as well as astrocytic markers (marked by *GFAP*, 68 nuclei). The neutrophil and ambiguous clusters were discarded, leaving only microglia and brain macrophages for downstream analysis (Supplementary Fig. 1c). Cluster markers are provided in Supplementary Table 6.

In order to determine if nuclei could recover microglial clusters identified in cells, we selected the top 40 markers defined by Mancuso *et al.* (Mancuso et al., 2019) for each of the clusters they identified in the original analysis of microglial cells. For each nucleus, we scored each set of markers based on the abundance of those markers in the nucleus, using Seurat’s *AddModuleScore* function. These scores, referred to as MS40 scores, are highlighted in Supplementary Fig. 1d. Our nuclei were able to recover a cytokine response cluster (CRM), marked by *CCL3*, *CCL4*, and an activation-like cluster, equivalent to the “*in vitro* microglia” identified in the original study (original markers included *APOC1*, *GPNMB*, *SPP1*, *APOE*). Homeostatic markers appeared ubiquitously through-out the nuclei dataset, and we were not able to distinguish a reduction of these markers in the activation-like response cluster, as we would expect from transcriptomic profiling of microglia in mice (Keren-Shaul et al., 2017; Sala Frigerio et al., 2019). Finally, the CAM (macrophage) cluster (*CD163*, *MRC1*), separated out from the bulk of the microglia, and was easily-recognisable based on its MS40 score. Cluster markers are provided in Supplementary Table 6.

In order to quantify the differences between cells and nuclei in more detail, we examined the overlap of the top 40 markers between nuclei clusters and cell clusters (Supplementary Fig. 1e). The cell macrophage (CAM) and cell cytokine (CRM) clusters showed the largest overlaps with Nuc1 and Nuc7 (27 and 24 of 40 markers, respectively). Other clusters only showed overlaps of between 1 and 5 genes. Cluster Nuc3 showed similar overlaps between “*in vitro* 1” and “*in vitro* 2” (5 genes). Cluster Nuc0 showed an overlap of 5 genes with “*in vivo* HM”, and cluster Nuc2 showed an overlap of 2 genes with “*in vivo* HM”. Cluster Nuc4 showed similarities with the “*in vitro* 2” cluster, suggesting it could be a cluster of activation, however all 5 overlapping genes were mitochondrial genes. Cluster Nuc3 markers *RPS12*, *TPT1*, *FTL*, *RPS18* and *EEF1A1* also appeared as markers of “*in vitro* 2”.

We performed similar analyses using more markers, however we found that introducing more markers resulted in nuclei markers overlapping with more than one cell cluster. We also noticed that introducing more markers resulted in overlaps between markers of the cellular clusters with each other. Selecting 40 markers allowed us to align cellular and nuclear clusters in an almost one-to-one fashion (see Supplementary Fig. 1e).

Overall, cytokine clusters and macrophage clusters were recovered well using single nucleus methods, however, differences between other microglial subpopulations were not convincingly recovered.

